# Comparative single cell analysis of wound and cancer identifies the metabolic dialogues between tumor initiating stem cells and macrophages

**DOI:** 10.64898/2026.04.02.716187

**Authors:** Weijie Guo, Daniel Leon, Benjamin Nicholson, Jianwen Que, Yuxuan Miao

**Affiliations:** Ben May Department of Cancer Research, The University of Chicago, Chicago, IL, USA; Columbia Center for Human Development and Division of Digestive and Liver Disease, Department of Medicine, Vagelos College of Physicians and Surgeons, Columbia University Irving Medical Center, New York, NY, USA

## Abstract

Macrophages are pivotal mediators of wound healing, yet the cellular programs they employ can be hijacked by cancers to drive tumorigenesis. Although similar macrophage programs support both physiological tissue regeneration and pathological cell growth, the molecular and functional difference between wound-associated macrophages (WAMs) and tumor-associated macrophages (TAMs) remain poorly defined. Here, we perform comparative single-cell RNA sequencing to delineate the dynamic cell states of macrophages during skin wound healing and the progression of cutaneous squamous cell carcinoma. Our analyses reveal that aberrantly regulated lipid metabolism is a distinct feature of TAMs. Critically, our genetic manipulations allow us to identify SOX2^High^ tumor-initiating stem cells as key orchestrators that modulate the lipid metabolism of TAMs and shape their cell states. These findings suggest that disrupting the metabolic crosstalk between tumor-initiating stem cells and TAMs represents a promising strategy to normalize myeloid cell function and enhance cancer immunotherapy efficacy.

## Introduction

Macrophages are highly plastic myeloid cells that can rapidly shift their phenotypes and cell states in response to diverse environmental cues (Locati et al., 2020; Wynn et al., 2013). Stimulation by pathogen- or damage-associated molecular patterns drives macrophages toward pro-inflammatory states, whereas exposure to immune-modulatory ligands promotes alternative activation and regulatory functions (Biswas and Mantovani, 2010; Mosser and Edwards, 2008; Murray et al., 2014). Their versatility and plasticity enable macrophages to play central roles in facilitating tissue repair (Vannella and Wynn, 2017; Wynn and Vannella, 2016). Immediately following injury, tissue-resident and monocyte-derived macrophages adopt pro-inflammatory functional states to control infection and clear cellular debris (Wynn and Vannella, 2016). As repair progresses to the healing phase, macrophages promptly transition to pro-healing states to suppress inflammation and induce angiogenesis (Shook et al., 2016; Shook et al., 2018). Given that stem cells are the principal mediators of tissue regeneration, a critical function of wound-associated macrophages (WAMs) during wound repair is to support stem cell self-renewal and differentiation, creating a niche that enables stem cells to efficiently restore tissue integrity (Shook et al., 2016; Shook et al., 2018).

Although many macrophage functions are essential for normal tissue repair, these programs are frequently co-opted by tumor-associated macrophages (TAMs) to promote tumorigenesis. As one of the most abundant immune cell populations in the tumor microenvironment (TME), TAMs drive cancer cell proliferation, enhance angiogenesis, and most importantly, establish an immunosuppressive milieu (Biswas and Mantovani, 2010). Indeed, similar myeloid cell responses underpin the conceptualization of cancer as “a wound that never heals” (Hu et al., 2023). Supporting this notion, recent spatiotemporal transcriptomic mapping of macrophage states during wound healing has highlighted processes such as antigen presentation and interferon responses are similarly activated in WAMs and TAMs (Hu et al., 2023). Notably, both WAMs and TAMs are thought to adopt the alternative activation (or M2-like) state, facilitating direct interactions with stem cells. It is increasingly evident that many solid tumors are initiated and maintained by a subset of tumor-initiating stem cells (tSCs) characterized by robust stemness signatures, akin to those responsible for normal tissue maintenance and repair (Ricci-Vitiani et al., 2007; Singh et al., 2004). These tSCs activate unique molecular programs to drive tumorigenesis and therapy resistance. In particular, they can survive robust immunotherapy treatments which otherwise effectively eradicates differentiated cancer cells (Miao et al., 2019; Oshimori et al., 2015). Similar to normal tissue regenerations, close interactions with the microenvironment are essential for tSCs to gain extrinsic protections during tumorigenesis (Guo et al., 2017; Lu et al., 2014; Taniguchi et al., 2020). Imaging analyses in skin squamous cell carcinomas (SCCs) have revealed that a subset of FcεRI+ macrophages cluster near tSCs (Casanova-Acebes et al., 2021), and these TAMs can produce signaling molecules—such as Wnt and TGFβ ligands—that activate corresponding pathways in tSCs (Oshimori et al., 2015; Taniguchi et al., 2020). Deciphering the reciprocal interactions between stem cells and macrophages is crucial for preventing cancer progression.

While previous research has largely focused on pathways that are similarly activated in macrophages during wound healing and tumorigenesis, the precise degree of similarity in cell states between WAMs and TAMs—along with any molecular features uniquely activated in either group—has not been comprehensively assessed. A fundamental difference between wound healing and cancer is that, whereas regeneration can conclude with resolution of inflammation and restoration of tissue architecture, these processes persist in cancers (Ge and Fuchs, 2018; Ge et al., 2017). Consequently, macrophages in tumors are chronically reeducated by constant cytokine signaling and reprogrammed by aberrant metabolic cues within the TME, endowing them with additional properties (Casanova-Acebes et al., 2021; Franklin et al., 2014; Hegde et al., 2025). Identifying the unique features of WAMs and TAMs and elucidating the respective mechanisms shaping their distinct cell states, may hold the key for developing strategies that normalize macrophage function in the TME as a novel approach to cancer immunotherapy.

In this study, we employed a partial-thickness wounding model to characterize the molecular features of macrophages that promote wound repair (Ge et al., 2017). This unconventional skin injury model removes only the superficial epidermis and upper hair follicle (Luan et al., 2024). In contrast to classical full-thickness wounds which are repaired by collective migration and proliferation of epidermal keratinocytes within a migrating tongue, in partial-thickness wounds, individual hair follicle stem cells (HFSCs) migrate into the wound bed (Luan et al., 2024). Our previous work demonstrated that, unlike full-thickness punch biopsy wounds where a migrating tongue shields underlying stem cells, partial-thickness wounds expose activated HFSCs directly to inflammatory insults, yet these stem cells can still be repurposed from hair regeneration to epidermis regeneration (Gonzales et al., 2021). This study suggests that stem cell survival requires temporary niche protection, potentially provided by macrophages with anti-inflammatory functions. In parallel, we investigated tumor-promoting macrophages in a chemically induced, autochthonous mouse model of cutaneous SCC, generated by sequential treatment with 7,12-dimethylbenz[a]anthracene (DMBA) and phorbol ester 12-O-tetradecanoylphorbol 13-acetate (TPA) (Nassar et al., 2015; Quintanilla et al., 1986). This cancer model enables the development of spontaneous SCCs with heterogeneous cell populations and hierarchical developmental trajectory, better recapitulating human epithelial cancers than transplanted models (Guo W, 2025). Importantly, SCCs in this system arise from mutated keratinocytes(Lapouge et al., 2011)—the same epithelial stem cells driving the wound repair—thus allowing direct comparison of macrophage signatures driving physiological versus pathological growth of the same tissue (Huang et al., 2017).

Using these refined models, we performed comparative single-cell analysis of dynamic macrophage states during skin wound healing and cutaneous malignant progression. Through transcriptome comparisons of WAMs and TAMs, we further characterized changes in macrophage heterogeneity and cell states in response to immunotherapy, with and without perturbation of critical pathways in tSCs. This analysis identified a pivotal role for a subset of SOX2^High^ tSCs in sculpting the unique phenotypes of a subset of TAMs, enabling these specialized TAMs to persist through immunotherapy and reciprocally protect tSCs.

## Results

### Global overview of the immune landscape in wound and cancer

To compare the molecular features of immune responses elicited during physiological tissue regeneration (e.g., wound healing) and pathological cell growth (e.g., cancer), we subjected immune cells from different stages of partial-thickness skin wounds, as well as various phases of skin squamous cell carcinoma (SCC) progression to single-cell RNA sequencing (scRNA-seq) (Fig. 1a). Specifically, we used fluorescence-activated cell sorting (FACS) to isolate CD45^+^ immune cells from Day 2 (inflammatory stage) and Day 4 (healing stage) wounds in mice, and from mice bearing skin papilloma (benign) or carcinoma (malignant) (two replicates per group; two pooled mice or tumors per replicate) (Fig. 1a). These timepoints were chosen based on previous studies characterizing the kinetics of partial thickness wound healing (Ge et al., 2017; Gonzales et al., 2021; Luan et al., 2024). Staging of SCCs was determined by their distinct appearance, necrotic states, tumor size, and histology. To profile transcriptomic similarities and differences among immune populations under these conditions, we employed a split-pool combinatorial barcoding-based scRNA-seq approach (Rosenberg et al., 2018). This method allows for the fixation, labeling, pooling, and simultaneous sequencing of cells from various conditions and time points, thereby minimizing batch effects (Rosenberg et al., 2018).

**Figure 1.**
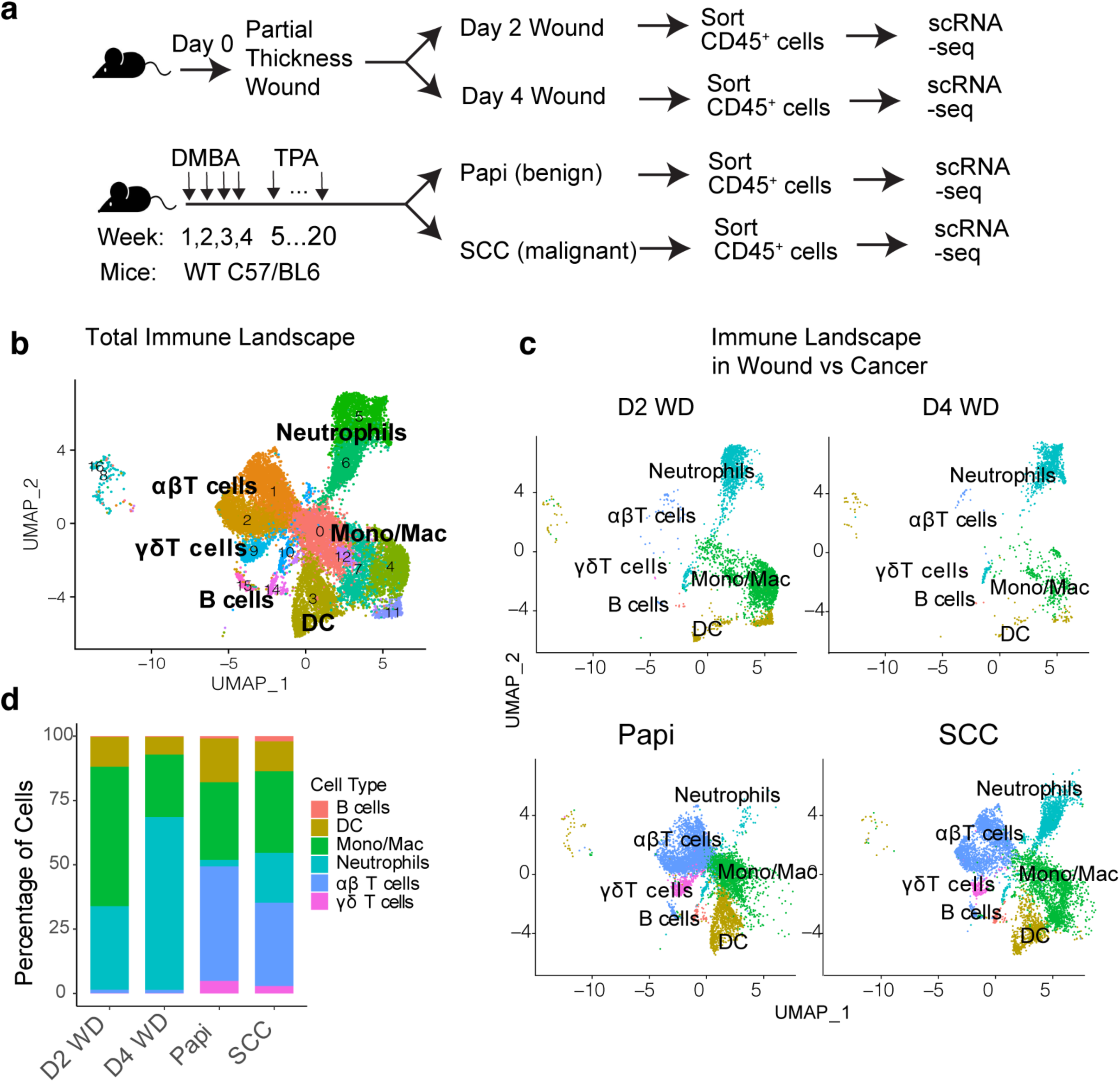
Global immune landscape in skin wound and cancer. **a.** Schematic of experimental procedures for inducing partial thickness wound and spontaneous skin SCCs for analyzing the immune landscape changes. **b.** UMAP showing the immune cell types identified by scRNA-seq. **c.** UMAP showing the representation of each major immune population identified in different stages of wound repair and skin cancer progression. **d.** Stacked bar chart showing the changes in the composition of immune landscape.

Following sequencing and quality control, we selected 18,456 immune cells for further analysis (3,781 cells from D2 WD, 1,492 cells from D4 WD, 6,601 cells from papilloma, and 6,582 from SCCs). Based on transcriptional profiles, we first examined major immune cell types and global changes in the immune landscape during wound healing and tumorigenesis. In total, we identified 17 distinct clusters when combining immune cells from both wounds and cancers (Fig. 1b). Among these clusters, we detected 10 clusters in the myeloid cell compartment and 7 clusters of lymphocytes (Fig. 1b). A global comparison of immune landscapes in wounds and cancers revealed key differences and similarities in the composition of these compartments. As expected, lymphocyte infiltration is the major difference between cancer and wound, and both T cells and B cells are mostly enriched in tumor samples (Fig. 1, c and d, Fig. S1a). A particularly notable T cell population that influxes into the tumor is the Foxp3^+^ regulatory T (Treg) cells (Fig. S1b). Although Treg cells have been shown to also play a key roles in wound repair, the presence of Treg in tumor samples drastically overshadows the wounded tissue, counteracting the anti-tumor functions of CD8^+^ T cells (Fig. S1b). In contrast, myeloid cells dominated the CD45^+^ immune cells in both wounds and cancers, accounting for more than 90% of the immune population in wounds and over 60% in cancers (Fig. 1, c and d). While the presence of neutrophils is comparable between wound and cancer (Fig. S1c), the dendritic cells are mostly present in cancer tissue (Fig. S1d). Overall, our data provided an important resource for the field to better understand the immune responses in both physiological tissue regeneration and pathological cell growth.

Given the critical roles of macrophages in both wound healing and cancer progression, in this study, we specifically focused on comprehensive comparison of the molecular signatures of these myeloid cells when they were driving wound repair and tumorigenesis. From our scRNA-seq data, we identified 7 major cell clusters of macrophages (Fig. 2a). Each cluster is enriched with markers (Fig. 2b) which can define the unique molecular features and distinct cell states. For example, the cells in cluster 0 (C0) are M1-like, being mostly enriched with proinflammatory genes, such as genes involved in antigen presentation and providing co-stimulatory signals (Fig. 2c), whereas the cells in cluster 1 (C1) were mostly enriched with signature genes that are known to define M2-like macrophages, such as *Mrc1* (genes encoding CD206), *Arg1* and *Cd68* (Fig. 2c). Compared to these two clusters, most macrophages found in the wound or cancer are in a “hybrid” cell state, being simultaneously enriched with both M1 and M2-like signatures (Fig. 2c). This dataset allowed us to identify the unique cell states that are associated with either wound healing or cancer progression.

**Figure 2.**
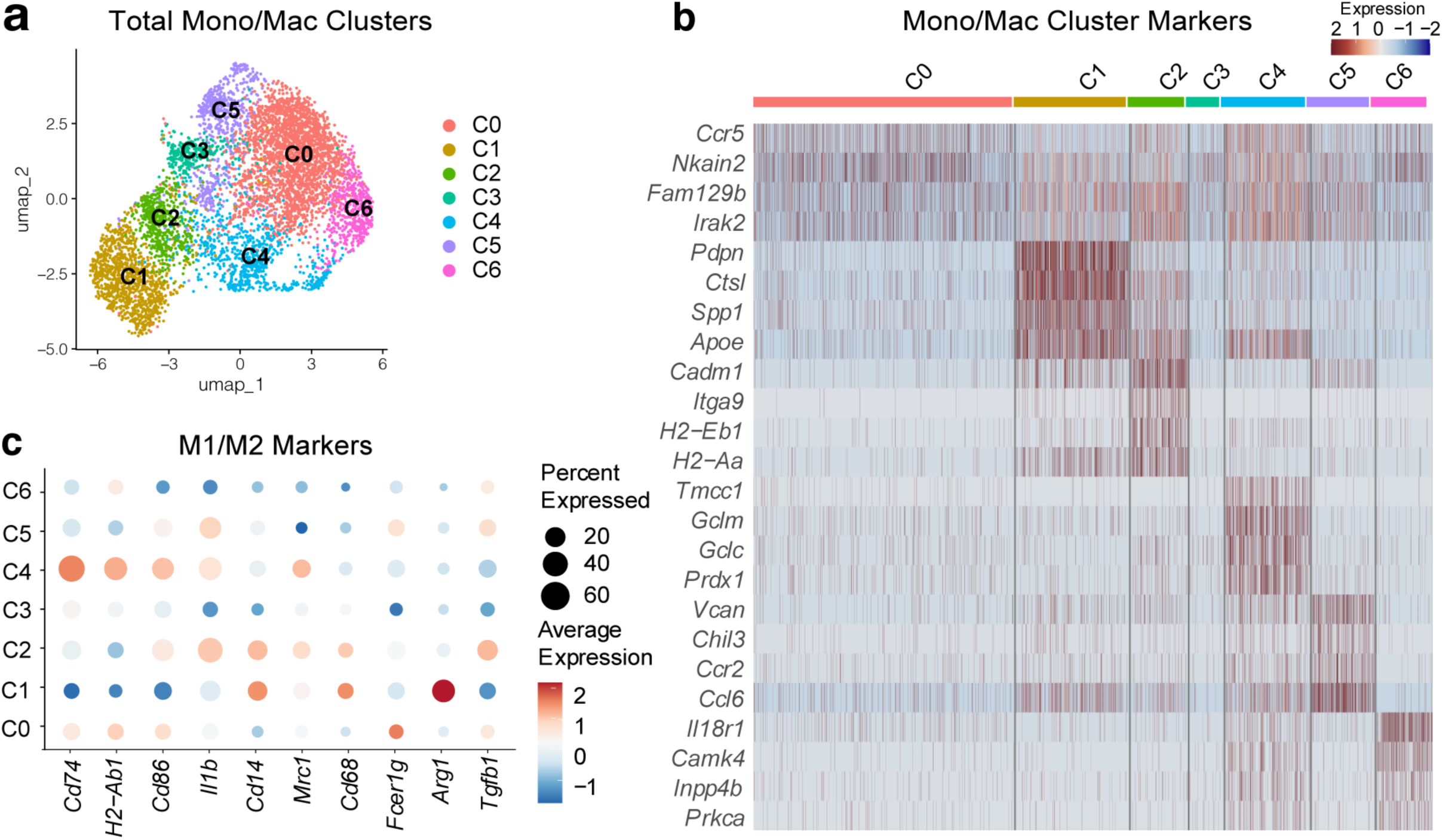
Global molecular features of macrophages during wound repair and cancer progression. **a.** UMAP showing the major cell states of macrophage identified by scRNA-seq. **b.** Heatmap showing the markers enriched in each identified macrophage cluster. **c.** Bubble heatmap showing the expression pattern of markers that are associated with either pro-inflammatory (M1-like) or anti-inflammatory (M2-like) in each identified macrophage cluster.

### Respective molecular features of macrophages during wound repair and tumorigenesis

We first focused on the dynamic cell states of macrophages during different stages of wound repair. Among all the macrophage clusters identified from our single cell analysis, we found four major clusters (C0, C1, C2, and C3) populating the wounded skin (Fig. 3a). We next sought to gain an in depth understanding into the molecular features of each major WAM subpopulations. To achieve this goal, we performed pseudo-bulk analysis of each cluster, followed by differential gene expression analysis of each cluster. Based on previous phenotype characterizations, it was established that distinct macrophage functional states drive different stages of wound healing. But the previous research has assumed an over-simplified paradigm. For example, the pro-inflammatory macrophages are thought to be the first responders being recruited immediately upon wounding to clear the dead cells or microbes (Mosser and Edwards, 2008; Wynn and Vannella, 2016). When the wound healing progress to the regenerative phase after the stem cells have migrated into the wound bed, these proinflammatory macrophages are converted into anti-inflammatory macrophages to promote re-epithelization and tissue remodeling (Mosser and Edwards, 2008; Wynn and Vannella, 2016). Consistent with this paradigm, our analysis confirmed that the cluster 0 (C0) WAMs which are mainly enriched with pro-inflammatory gene signatures were mostly found in Day 2 post-wounding, and by Day 4, majority of these pro-inflammatory macrophages were eliminated from the wounded tissue (Fig. 3, a and b). Even with a few C0 cells remaining in the wounded skin by Day 4, these cells turned off their pro-inflammatory programs (Fig. 3c). Unlike the pattern of proinflammatory macrophages which was consistent with previously reported, to our surprise, the emergence of pro-healing macrophages was fairly early. Although the hair follicle stem cells won’t migrate into the damaged epidermis until Day3 (Ge et al., 2017; Gonzales et al., 2021; Luan et al., 2024), as early as Day 2 post-wounding, more than 60% of WAMs were already in the cell states which are enriched with genes involved in immune suppression, lipid metabolism, and tissue regeneration (Fig. 3, c to e). Another interesting finding from this analysis was that, in contrast to the pro-inflammatory macrophages that had limited heterogeneity (mainly in C0 cluster), we found three distinct clusters of macrophages harboring the potential of promoting tissue regeneration and suppressing inflammation (Fig. 3, a and b). Importantly, each of these three clusters activated a different set of genes that could contribute to different aspects of tissue regeneration. For instance, among all the macrophages, the C1 cells were defined as their highest level of *Arg1* (Fig. 2c) and *Ptges* (Fig. 3e), while the cells in C2 expressed highest level of *Tgfβ1* (Fig. 2c) and *Il10* (Fig. 3c). Both C1 and C2 macrophages also blunted the production of pro-inflammatory chemokines when the wound healing progressed from Day 2 to Day 4 (Fig. 3c). Similarly, C1 WAMs were the major population expressing *Ereg* and *Pf4*, and C2 WAMs expressed *Chil3* (Fig. 3e). With these coordinated actions, different subsets of WAMs functioned synergistically to dampen down the inflammation and promote stem cell-mediated re-epithelization and differentiation. Interestingly, compared to the proinflammatory C0 WAMs, another distinct molecular feature of WAMs in both C1 and C2 is their enrichment of genes involved in lipid metabolism, such as *Spp1, Lpl, Cd36, Fabp5,* and *Apoe* (Fig. 3d). These genes were particularly high in C1 WAMs, and these genes were already upregulated on Day 2 post-wounding (Fig. 3e). These findings suggested that the elevated lipid metabolism is a unique requirement for macrophage to promote tissue growth.

**Figure 3.**
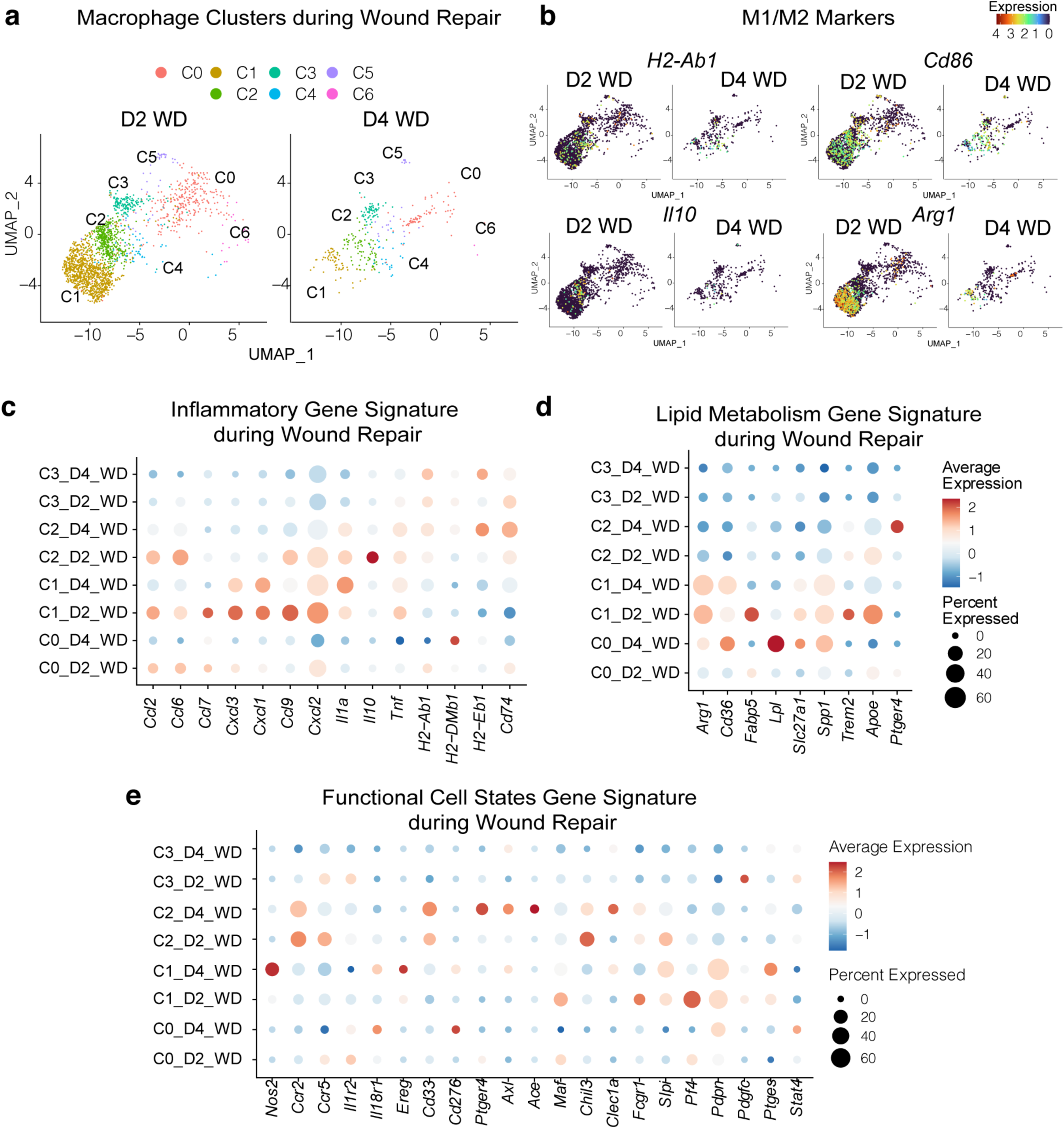
Dynamic molecular features of macrophages during wound repair. **a.** UMAP showing the dynamic changes of each major macrophage subpopulation during different stages of wound repair. **b.** UMAP showing the expression pattern of pro-inflammatory (*H2-Ab1* or *Cd86*) or anti-inflammatory (*Il10* or *Arg1*) genes in major macrophage subpopulations present in different stages of wound repair. **c** to **e**. Bubble heatmap showing the transcripts of genes encoding important factors promoting (**c**) inflammation, (**d**) lipid metabolism, and (**e**) various functional states of macrophages during different stages of wound repair.

Next, we sought to investigate the evolving features of TAMs when DMBA-induced skin tumors progress from benign (papilloma or papi) to malignant stage (SCC). Four distinct clusters (C0, C4, C5, and C6) of macrophages were identified in the tumors (Fig. 4a). When we closely examined the representation of these clusters in different stages of tumorigenesis, we first found that the benign tumors are mostly enriched with C0 TAMs that activate strong antigen-presentation abilities (Fig. 4, a and b) and the C6 TAMs that express high level of interferon stimulated genes (Fig. 4, a and c). When the cancer progressed from benign to malignant stage, these anti-tumor TAMs were significantly reduced (Fig. 4a). In each cluster, the expression level of genes encoding the antigen presenting machinery (Fig. 4b) and providing interferon responsive potential (e.g. *Ifngr1*) were significant reduced (Fig. 4c), while the genes that can repress the interferon response (e.g. *Ptpn7*) increased (Fig. 4c). This was consistent with the established idea that the host immunity can still control the benign tumors, whereas the establishment of the immune suppression by the remodeling of the TME was the prerequisite of the malignant progression. In parallel with the contraction and functional remodeling of the anti-tumor macrophage compartments, several new macrophage cell states (C4 and C5) emerged at the malignant SCC stage (Fig. 4a). Most genes that were uniquely activated in these two new cell states of TAMs during the SCC stage were involved in suppressing anti-tumor immunity (e.g. *Cd274, Mmp12*), escalating pathological inflammation (e.g. *Il1a, C1qa*), and promoting cell growth (e.g. *Pdgf*) (Fig. 4, d and e). Among all the molecular features acquired by the C4 and C5 clusters of pro-tumor TAMs newly emerged during malignant progression, the enrichment of lipid metabolism-related genes, such as *ApoE and Cd36*, were particularly interesting (Fig. 4f), especially in C5 TAMs. The similar activation of lipid metabolism related genes in both WAMs and TAMs further highlighted the important roles of these pathways and processes for macrophages to promote regeneration and growth. Taken together, our results provide a comprehensive and dynamic overview of the evolving molecular features of TAMs during the malignant progression.

**Figure 4.**
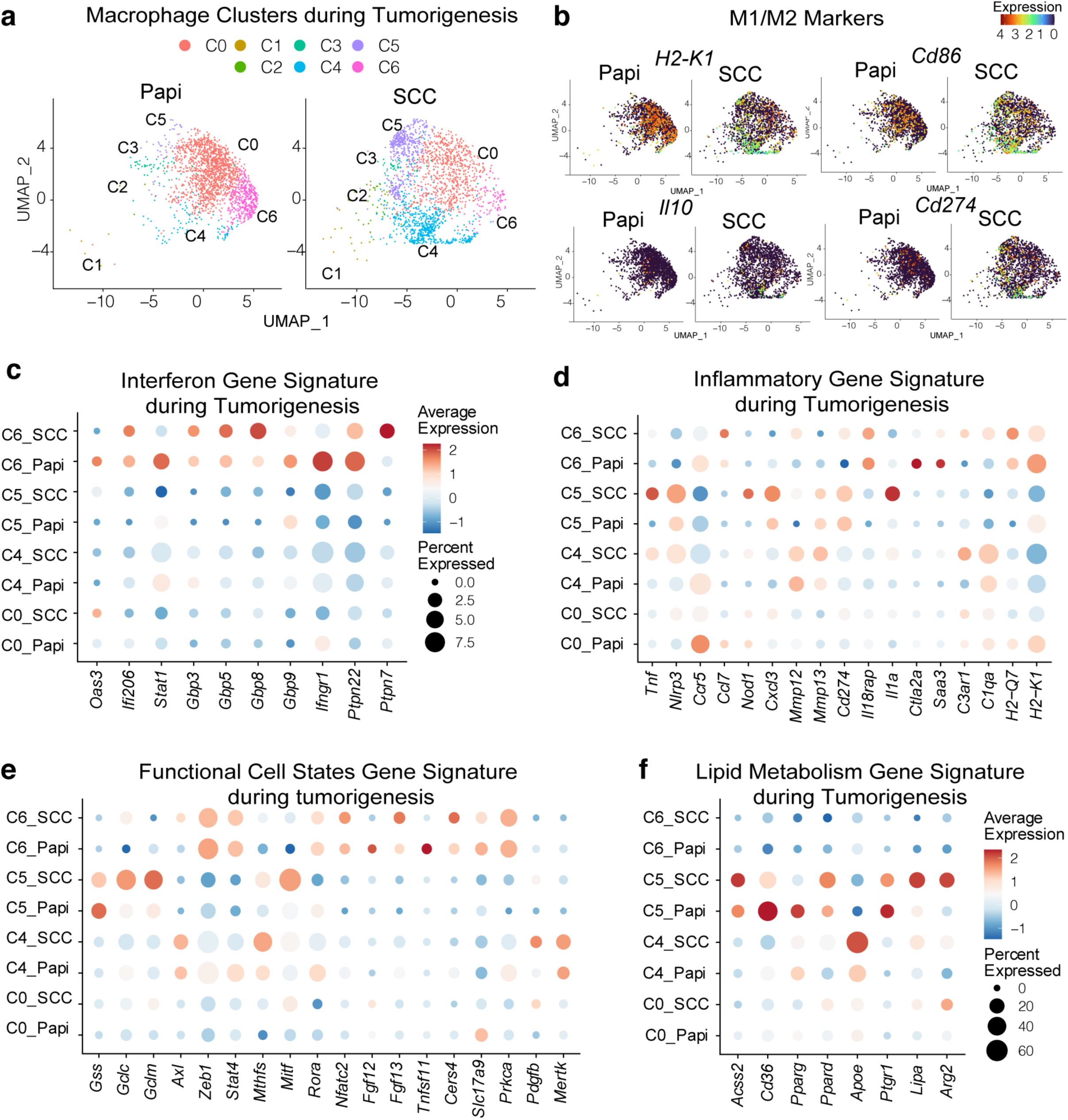
Evolving molecular features of macrophages during cancer progression. **a.** UMAP showing the evolving changes of each major macrophage subpopulation during different stages of skin cancer progression. **b.** UMAP showing the expression pattern of pro-inflammatory (*H2-Ab1* or *Cd86*) or anti-inflammatory (*Il10* or *Cd274*) genes in major macrophage subpopulations present in different stages of skin cancer progression. **c** to **f**. Bubble heatmap showing the transcripts of genes encoding important factors promoting (**c**) interferon response, (**d**) inflammation, (**e**) various functions of macrophages, and (**f**) lipid metabolism during different stages of skin cancer progression.

### Comprehensive comparison of macrophage signatures in wound and cancer

Based on the insights gained from analyzing WAMs and TAMs separately, we next sought to compare their molecular features. To this end, we first computationally overlaid the clusters of WAMs and TAMs. Interestingly, although macrophages play similar roles in promoting cell growth and share many key functions during both wound repair and tumorigenesis, we found that the C0 cluster was the only cell state that was shared between skin wounds and skin cancers (Fig. 5, a and b). Consistent with previous reports(Hu et al., 2023), the antigen presentation and co-stimulation signature were indeed similarly activated in C0 cluster in both wound and cancer. (Fig. 3b and Fig. 4b). However, most other cell stats of WAMs and TAMs were highly distinct (Fig. 5a and 5b). As shown in Figure 5A and 5B, the C1, C2 and C3 were found mainly in wounded skin, while C4, C5, and C6 represented the cell states of macrophage in cancer. RNA velocity indicated the developmental trajectory of these macrophage clusters, showing that cell states identified in WAMs appear to be more intermediate on the trajectory, since these clusters can differentiate towards the C4 and C5 states found in TAMs (Fig. 5c).

**Figure 5.**
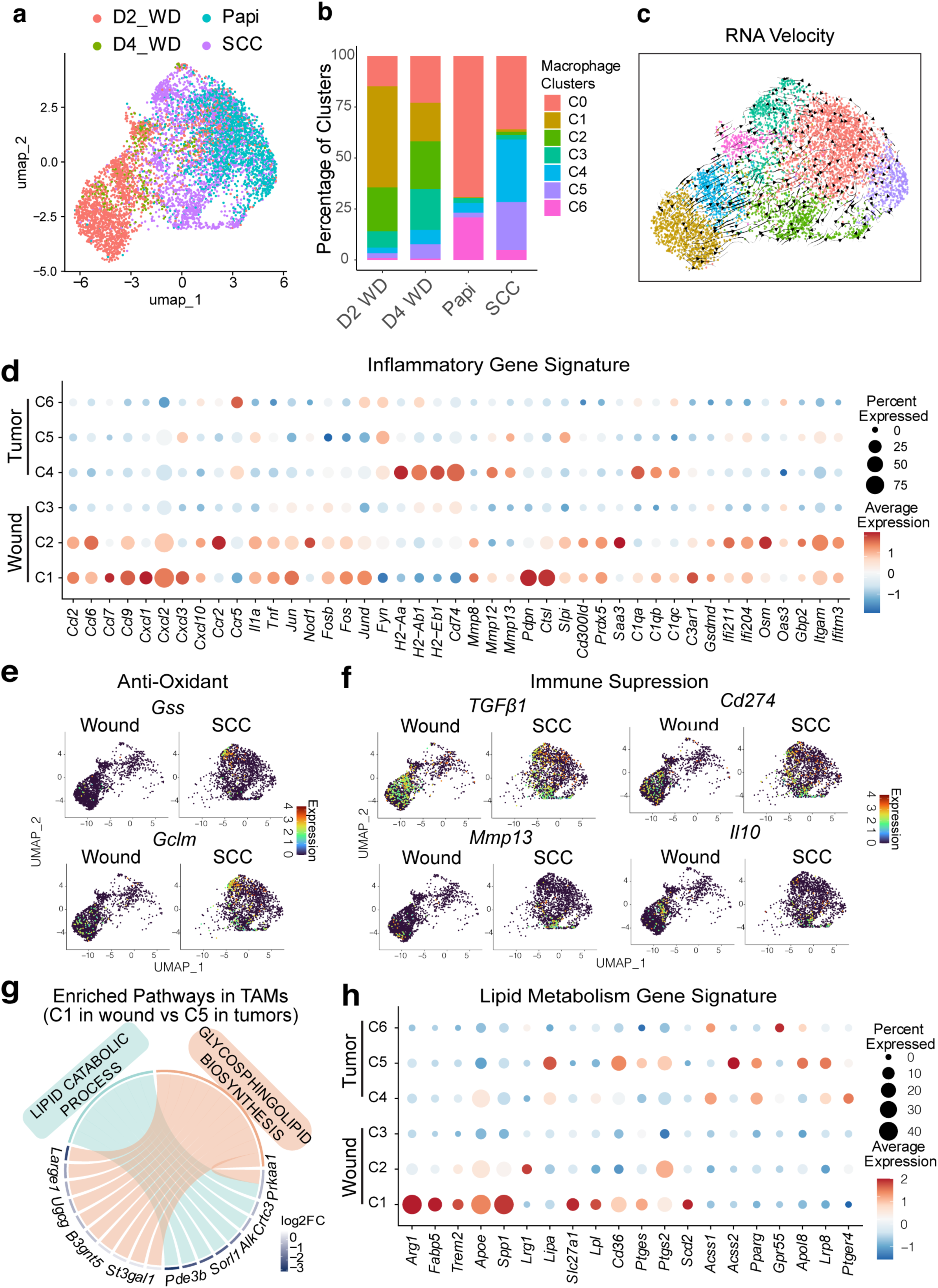
Comparative analysis of molecular signatures of macrophages in wound and cancer. **a.** UMAP showing the overlaid macrophage subpopulation found in wound and cancer. **b.** Stacked bar chart comparing the changes in the composition of macrophage population in wound and cancer. **c.** RNA velocity analysis of potential developmental trajectory and relationship among various macrophage clusters identified in wound and in cancers. **d.** Bubble heatmap showing the transcripts of inflammatory genes in the major macrophage clusters that are distinctively present in wound (C2, C3, and C4) and cancer (C4, C5 and C6). **e** and **f**. UMAP showing the expression pattern of (**e**) anti-oxidant (*Gss* or *Gclm*) or (**f**) immune suppression (*Tgfβ1, Cd274, Mmp13, or Il10*) genes in major macrophage subpopulations present in different stages of wound healing or skin cancer progression. **g.** Chord diagram showing the up-regulated lipid metabolism related pathways in the macrophages found skin SCC compared to macrophage found in skin wounds. **h.** Bubble heatmap showing the transcripts of genes encoding important factors promoting lipid metabolism during different stages of wound healing and skin SCC progression.

We next focused on the specific programs that can distinguish the WAMs and TAMs. To this end, we performed pseudo-bulk differential gene expression analysis. We specially focused on comparing the M2-like C1 and C2 clusters found in the wound with the similar anti-inflammatory C4 and C5 clusters that emerged when cancer progressed to carcinoma. We first found that both WAMs and TAMs displayed distinct inflammation-regulatory profiles. The complement-related genes, such as C1qa, C1qb and C1qc, were particularly upregulated in TAMs, whereas many chemokines (e.g. *Cxcl1, Cxcl3, Ccl2,* and *Ccl9*) and cytokines (e.g. *Tnf, Il1a*), were expressed at higher level in WAMs (Fig. 5d). Secondly, it was also intriguing that majority of TAMs emerged during the SCC stage were enriched with the antioxidant and express higher level of immune evasive genes (Fig. 5, e and f). This finding suggests that unique molecular adaptations were necessary for TAM establishment within the tumor microenvironment. To our surprise, although the anti-inflammatory macrophages in both wound and cancer were enriched with lipid metabolism signatures (Fig. 3d and Fig. 4f), additional regulatory mechanism appeared to be only activated in TAMs to further reprogram their lipid metabolism (Fig. 5, g and h). Specifically, although genes, such as *Trem2, ApoE, Lpl, and Cd36,* were upregulated in both C1/C2 of WAMs and C4/C5 of TAMs, additional receptors (e.g. *Lrp8, Gpr55*), new transcription factor (e.g. *Pparg*), as well as new enzymes (e.g. *Acss1, Acss2*) were further upregulated in TAMs (Fig. 5h). This key finding suggested that the aberrant lipid metabolism was a defining feature that distinguished the functional cell states of macrophages during pathological growth after transformation from the physiological regeneration during wound repair. This finding inspired us to further explore the underlying mechanisms that are uniquely activated in cancer that induce the aberrant regulation specifically in TAMs.

### SOX2^High^ tSCs shaped the metabolic states of TAMs

Given the shared roles of macrophages in promoting rapid cell proliferation, establishing immune suppression, and facilitating tissue remodeling in both wound healing and cancer, it was striking to observe extensive distinctions in the macrophage cell states in wound and cancer. To dissect the underlying basis that shaped the aberrant cell states of TAMs, we focused on the factors and mechanisms that were uniquely activated in cancers. The specific expression of FcεR1 (Fig. 2c) in TAM subpopulations that are enriched with lipid metabolism related genes was particularly intriguing. Recent investigations have shed light into this question. FcεR1^+^ TAMs are predominantly located at the tumor-stroma interface (Taniguchi et al., 2020), where they interact with TGFβ-responding tSCs. These tSCs recruit TAMs via IL33 secretion(Taniguchi et al., 2020), and in turn, these FcεR1+ macrophages provide reciprocal regulation by producing ligands such as TGFβ.(Raghavan et al., 2019) While normal epithelial stem cells modulate immune cells to facilitate wound repair, tSCs exploit this interaction via additional oncogene-driven mechanisms to drive tumorigenesis. Many such new processes are now known to be mediated transcription factors, such as SOX2 (Boumahdi et al., 2014). SOX2 plays key roles in embryonic stem cells, especially in facilitating their stemness programs (Boumahdi et al., 2014). Interestingly, this key transcription factor is silenced in many adult tissue cells but becomes reactivated in transformed cancer cells (Bass et al., 2009; Vanner et al., 2014). Since SOX2 is a key driver that can distinguish the activity of normal skin stem cells and tSCs (Boumahdi et al., 2014), and because SOX2 is known to induce a unique lipid profile in cancer cells (Shen et al., 2021; Shen et al., 2025; Wang et al., 2025), we specifically focused on exploring its potential role in endowing the stem cell-like cancer cells to modulate macrophage cell states through regulating their lipid metabolism.

To test this hypothesis, we developed two complementary genetic models. Since *Sox2* is silenced in normal skin epithelial stem cells, we first sought to ectopically express *Sox2* in skin epithelium and then examine how this genetic manipulation of normal epithelium impacts the cell states of WAMs in the wounds. For this purpose, we generated Sox2 conditional overexpression (cOE) mice (*K14CreER; R26-LSL-Sox2-IRES-GFP*) (Fig. 6a). Upon tamoxifen injection, we can activate Sox2 in basal skin epithelium where epidermal stem cells and hair follicle stem cells are located (Fig. 6a). In parallel, we also generated the *Sox2* conditional knockout (cKO) mice (*K14CreER; Sox2 ^flox/flox^*) (Fig. 6b). We again employed DMBA/TPA treatment to induce autochthonous skin SCCs on both Cre negative control and *Sox2* cKO mice, so we can profile the molecular changes in macrophages when *Sox2* is ablated in tSCs in a more physiologically relevant TME (Fig. 6b). Following tamoxifen treatment to induce either *Sox2* activation in normal SCs or *Sox2* ablation in tSCs, we isolated total CD45^+^ immune cells from either the wounds or from the SCCs, and subjected these cells to scRNA-seq (Fig. 6, a and b). In total, we again identified six major clusters of macrophages in both skin wounds and skin SCCs (Fig. 6c). Importantly, the cluster 2 (C2) contained the lipid laden macrophages that were enriched with the genes involved in lipid metabolism, such as *Spp1, Lpl* and *Cd36* (Fig. 6d).

**Figure 6.**
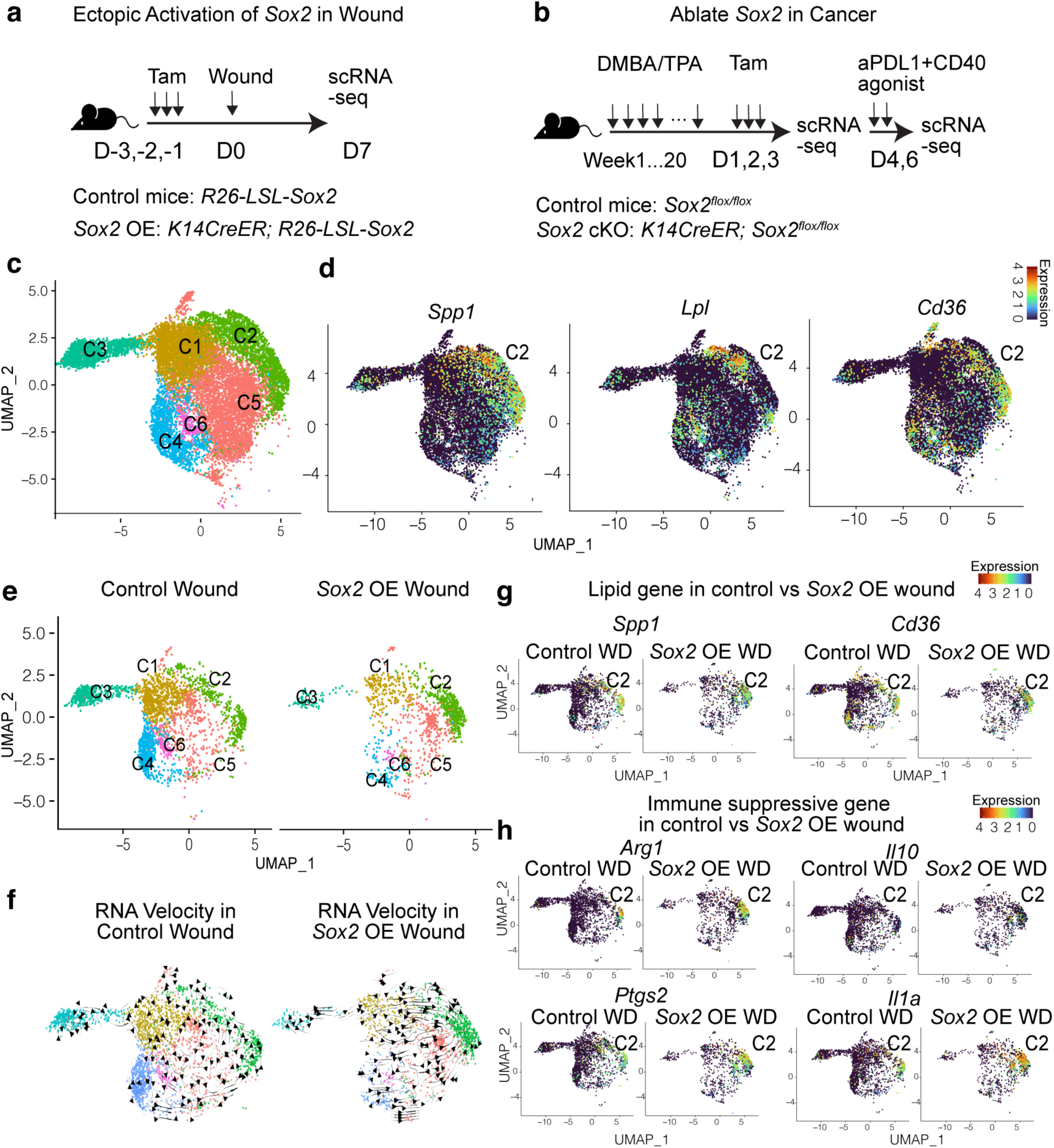
Ectopic activation of *Sox2* in skin epithelium during wound repair induces WAMs to acquire TAM-like signatures. **a** and **b**. Schematics showing the model for (**a**) ectopic activation of *Sox2* in normal skin epithelium during wound repair or (**b**) genetic ablation of Sox2 in tSCs in spontaneous skin SCCs before the anti-PDL1 + CD40 agonist immunotherapy treatment. **c.** UMAP showing the composition of macrophage subpopulations in skin SCC. **d.** UMAP showing expression of various lipid metabolism related genes specifically in the cluster 2 in TAMs. **e.** UMAP showing the expansion of C2 WAM, but shrinkage of other macrophage subpopulation when Sox2 is ectopically activated in normal skin epithelium during wound repair. **f.** RNA velocity analysis showing the developmental trajectory of other macrophage clusters towards the C2 WAMs when *Sox2* is ectopically activated in normal skin epithelium during wound repair. **g** and **h**. UMAP showing the upregulation of genes involved in (**g**) lipid metabolism and (**h**) immune suppression in C2 WAMs when *Sox2* is ectopically activated in normal skin epithelium during wound repair.

With this new dataset after various genetic manipulations, we first compared the macrophage landscape in control and *Sox2* cOE wounds to determine how ectopic activation of *Sox2* in skin epithelium impact macrophages. In the control wounds, as observed before, multiple cluster of WAMs with distinct molecular features co-exist to drive different processes of wound repair (Fig. 6e). However, when the *Sox2* is ectopically expressed in skin epithelium, at the same time point post wounding, most macrophages clusters shrink whereas we only observed significant expansion of C2 WAMs that are enriched with lipid metabolism genes (Fig. 6e). Interestingly, the RNA Velocity analysis shows that the developmental trajectories of macrophage cell states are distinct between control and *Sox2* cOE wounds (Fig. 6f). In contrast to the WAMs in control wounds, the other macrophage cell states differentiate towards the C2 WAMs in *Sox2* cOE wounds (Fig. 6f). When we compared the expression pattern of specific genes in the C2 cluster between control and *Sox2* cOE wounds, we find that most lipid metabolism-related genes, such as *Spp1* and *Cd36*, are upregulated in C2 WAMs in the *Sox2* cOE wounds (Fig. 6g). In consistent with the critical immune suppression and pro-tumor roles of this cluster cells, we also observed upregulation of genes involved in immune suppression, such as *Il10* and *Arg1*, and pathological inflammation, such as *Ptgs2* and *Il1a* (Fig. 6h) in *Sox2* cOE wounds. Taken together, these analysis allows us to confirm that, even without the oncogenic transformation and chronic inflammation exposure, when the epithelial stem cells start to express *Sox2*, the gene signatures of their surrounding WAMs start to resemble the lipid laden macrophage-like TAMs.

We next focused on the molecular changes in macrophages when *Sox2* is specifically ablated in K14^+^ basal epithelium in cancers. We have found that, after losing *Sox2* in cancer cells, majority of these lipid metabolism-related genes were ablated (Fig. 7, a and b) in the C2 TAMs. Instead, the same cluster of pro-tumor TAMs started to up-regulate genes known for mediating anti-tumor functions, such as *Nlrp3* (Han et al., 2021; Jing et al., 2023; Zhivaki and Kagan, 2021) and *Nos2 (Sun et al., 2021; Weiss et al., 2010)* (Fig. 7b), leading to significantly increase of T cells (Fig. 7c). These key results further confirmed our speculation that when the macrophages are recruited by tSCs (Taniguchi et al., 2020), these SOX2^High^ tSCs can also shape the cell states of macrophages, especially allowing these macrophages to acquire lipid laden macrophage-like phenotype that can distinguish the TAMs from WAMs.

**Figure 7.**
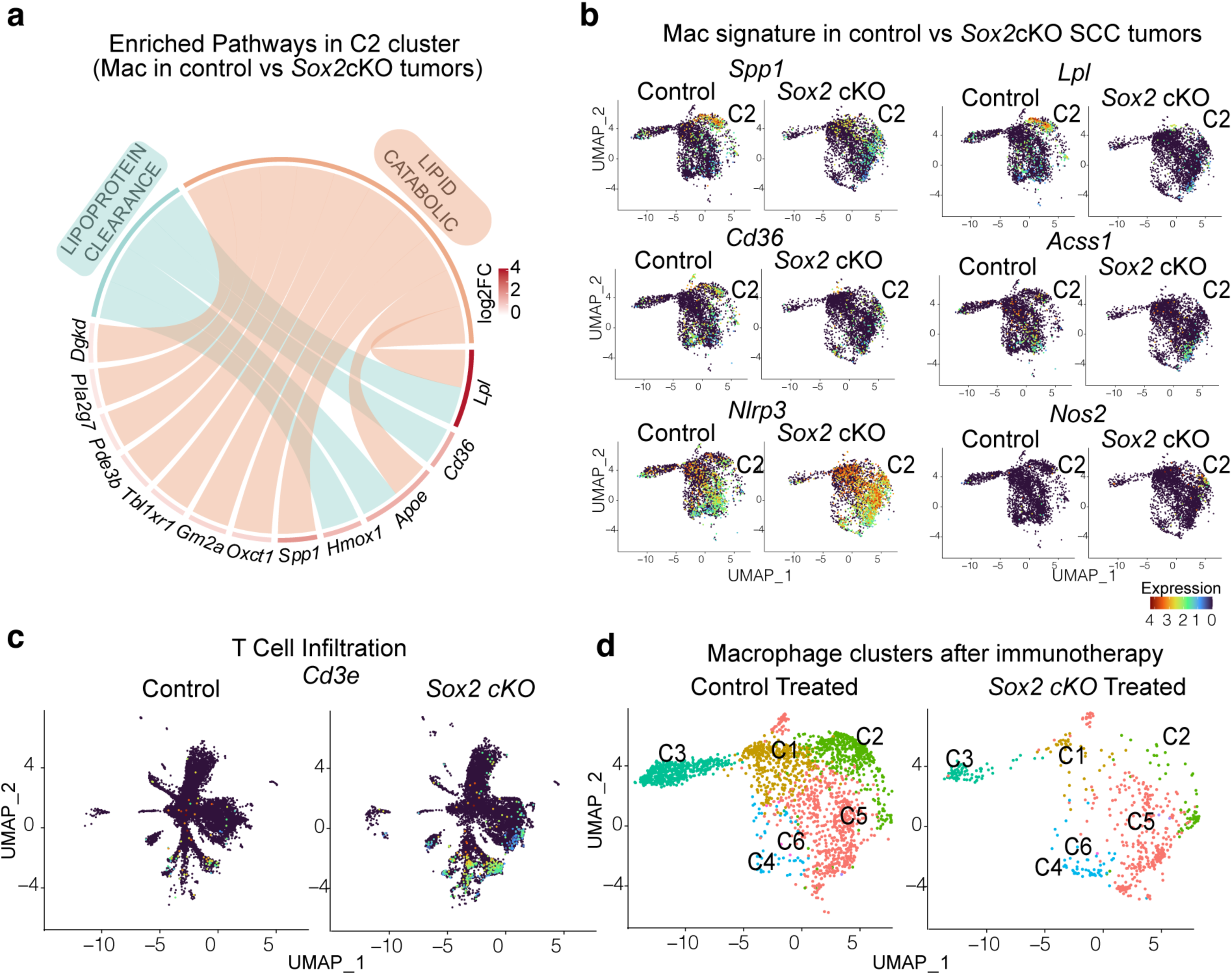
Genetic ablation of *Sox2* in tSCs blunt the lipid metabolism and maintenance of TAMs. **a.** Chord diagram showing the reduced enrichment of lipid metabolism related pathways in the macrophages found skin SCC after depletion *Sox2* in epithelial cancer cells, compared to macrophage found in control tumors. **b.** UMAP showing the expression of various lipid metabolism related genes specifically in the cluster 2 in TAMs with and without depleting *Sox2* in epithelial cancer cells. **c.** UMAP showing the increased infiltration of CD3^+^ T cells in skin SCCs with or without depleting *Sox2*. **d.** UMAP showing the composition of macrophage subpopulations in skin SCC with and without depleting *Sox2*.

### Metabolic crosstalk with tSCs was essential for TAMs to persist during immunotherapy treatment

To further determine the importance of the SOX2^High^ tSCs for modulating and maintaining the lipid laden TAMs, we treated both control and *Sox2* cKO mice bearing DMBA/TPA-induced spontaneous SCCs with anti-PDL1 + CD40 agonist immunotherapy. Recent study has shown that this combination of immunotherapy can induce robust anti-tumor immune response in cancers harboring high mutation burdens, such as skin SCC (Benguigui et al., 2024; Gungabeesoon et al., 2023). However, while the T cell activities can be reinvigorated to induce rapid tumor clearance, many treated tumors eventually grow back (Guo W, 2025). Our previous work demonstrated that the rapid tumor relapse is driven by the SOX2-mediated and stem cell-specific immune modulation, as the tumor relapse can be blocked when *Sox2* is ablated (Guo W, 2025). Here, we explored whether this effect of SOX2 was partially due to the loss of these lipid-laden macrophages. Single cell RNA-seq analysis first showed that, in the immunotherapy treated control tumors, many pro-tumor macrophages, especially the subset of *CD163*^+^ pro-tumor TAMs (shown in cluster 4 or C4), were replaced by pro-inflammatory macrophages with strong T cell priming activities (Fig. 7d). In contrast, most of the pro-tumor TAMs that were enriched with lipid metabolism-related genes, such as *Spp1, Lpl,* and *Cd36*, (shown in cluster 2 or C2) can persist (Fig. 7d). However, when the *Sox2* is ablated in the skin SCC epithelial cells, after immunotherapy treatment, these macrophages cannot persist in the TME anymore (Fig. 7d). Collectively, our study identifies SOX2^High^ tSCs as an unexpected regulatory cell population that emerges in cancer. These cells orchestrate the unique development and functional maintenance of immune-suppressive, lipid-laden TAMs, distinguishing them from the macrophages typically enriched in wound environments.

## Discussion

Macrophages play critical roles in both physiological tissue regeneration and malignant cell growth. While many studies highlighted important similarities between macrophages infiltrating wounded and cancerous tissue, the key distinctions and underlying regulatory mechanisms shaping the differences between these macrophages are still unexplored. In this study, by performing comparative single cell analysis of macrophages in skin cancers and skin wounds, we unveiled that the aberrant lipid metabolism is a previously unrecognized molecular feature of TAMs that might explain the unresolvable myeloid cell dysfunctions specifically found in cancers. Macrophages enriched for genes involved in lipid metabolism—commonly referred to as lipid-laden macrophages—have recently been identified as the predominant TAM population responsible for pro-tumor and immune suppressive functions (Governa et al., 2024; Kloosterman et al., 2024). In particular, as verified from our single cell analysis, these subsets of macrophage are the major subpopulation expressing many important TAM-associated markers, such as ARG1, SPP1, and TREM2 (Marelli et al., 2022). Thus, compared to many other macrophage subpopulations that have pro-regeneration functions in both injury repair and tumorigenesis, this subset of special TAMs might represent a special therapeutic target for normalizing the TME. Indeed, recent studies using TREM2 or SPP1 blocking antibodies has shown promising effects in treating several types of cancers in preclinical models (Kartal et al., 2025; Katzenelenbogen et al., 2020; Molgora et al., 2020; Sun et al., 2023). Thus, our study provided important explanations for these therapeutic effects of new treatments.

A particularly surprising finding from this study is that the development of these special macrophages appears to rely on a special subset of SOX2^high^ tumor initiating epithelial stem cells. Importantly, we have found that when we ectopically activated Sox2 expression in skin epithelium wound repair, most macrophage subpopulations are induced to differentiate towards macrophage cluster that are enriched in lipid metabolism and this cluster of cells also further upregulate their lipid metabolism, acquiring the gene signatures resembling the lipid laden TAMs. On the other hand, when we genetically ablated *Sox2* in cancer, these macrophages reduced their lipid metabolism, leaving them susceptible to the immunotherapy-induced reprogramming. It is still unclear how these SOX2^high^ tSCs modulate macrophages. Furthermore, the identity of the specific lipids or fatty acids produced by these tSCs to mediate this metabolic crosstalk represents a critical area for future investigation. Although these important questions are out of the scope and cannot be addressed in the current study, this study indeed unveiled an important strategy to design novel immunotherapies. Recent studies have demonstrated that these SOX2^high^ tSCs are the culprit of cancer relapse since they can survive from robust anti-tumor immune response. This study demonstrated that these tSCs may achieve this feature by modulating the differentiation of macrophages into lipid-laden macrophages, so they can form a special niche and provide reciprocal protections for these tSCs. Future investigation should focus on dissecting the molecular mechanisms employed by these SOX2^high^ tSCs to modulate macrophages. These future studies will uncover novel strategies to disrupt the tSC-macrophage communications, thereby removing this critical niche protection and sensitizing tSCs to T cell-mediated attack.

Taken together, this comprehensive comparative analysis provides an important resource for understanding the developmental trajectory and molecular features of tumor-associated macrophages. Moreover, it uncovers a key orchestrator that shapes the heterogeneity and functional states of TAMs.

## Materials and methods

### Mice and ethics

*K14CreER* mice were generated by Dr. Elaine Fuchs lab and backcrossed to C57/BL6J background for ten generations(Naik and Fuchs, 2022). Wild-type C57BL/6J, and *Sox2^flox/flox^ (B6.Sox2^tm1.1Lan^/J),* were obtained from The Jackson Laboratory. *LSL-Sox*2 mice were described before (Liu et al., 2013). To induce *Sox2* overexpression or conditional knockout, a daily i.p. injection of 2 mg tamoxifen for three consecutive days was performed.

All mice were maintained in an Association for Assessment and Accreditation of Laboratory Animal Care (AAALAC)-an accredited animal facility. Procedures were performed using IACUC-approved protocols. All mice were maintained and bred under specific-pathogen free conditions at the University of Chicago. All the procedures are in accordance with the Guide of the Care and Use of Laboratory Animals.

### Tumor formation and treatment

A previously established two-stage chemical carcinogenesis protocol was used to induce skin SCC formation. The shaved dorsal skin of various mouse lines was topically treated with 200 µL of 200 nM DMBA once a week for 4 weeks to initiate tumorigenesis. Four weeks later, tumor promotion was induced by weekly (twice a week) applications of 200 µL of 35 µM TPA for 20 weeks.

For immunotherapy treatment, tumor bearing mice were treated with a combination of anti-PD-L1 blocking antibody (Clone 10F.9G2, 100 µg/mouse, BioXCell) and anti-CD40 agonist antibody (Clone FGK4.5, 100 µg/mouse, BioXCell) intraperitoneal for a total of 2 doses every other day. Tumors were collected 2 days after the last treatment for downstream experiments.

### Partial thickness wound

The partial thickness wounding was performed following previously established protocols (Ge et al., 2017). Briefly, mice were shaved in second telogen (postnatal day P60). The remaining hair was removed using depilatory cream, and a Dremel drill head was used to gently scrape the back skin of anesthetized mice. To standardize the wound depth, the number of touches performed with the Dremel drill was pre-determined by inspecting the wounded skin for the first signs of erythema and pinpoint bleeding. This method removes the epidermis and upper HF, including most infundibulum and isthmus cells, but leaves the HF bulge intact.

### Cell sorting and flow cytometry

To sort immune cells from wounded skin, the wounds were collected at the selected time points and then placed in cold PBS for 15 minutes to remove the scab. The remainder of the wounded tissue was minced in digestion media [RPMI-1640 (GIBCO) with HEPES (25 mM, Corning), MEM non-essential amino acids (1:100, Thermo Fisher), solidum pyruvate (1 mM, Corning), penicillin (100 U/ml), streptomycin (100 μg /ml), gentamicin (20 μg/ml, Thermo Fisher), β-mercaptoethanol (55 μM, GIBCO)]. The minced tissue was digested with Liberase (25 μg/ml) (Roche) for 120 minutes at 37 °C with an adapted protocol (Luan et al., 2024). Single-cell suspensions were obtained, and cells were stained using an antibody cocktail prepared at predetermined concentrations in a staining buffer (PBS with 5% FBS, 5mM EDTA 25 mM HEPES).

To sort immune cells from DMBA/TPA induced tumors, the tumors were dissected at designated time points, minced and digested with Liberase (25 μg/ml) (Roche) in RPMI-1640 (GIBCO) for 60 minutes at 37 °C. Digested tissues were then passed through a 70 µm cell strainer, subjected to ACK lysis for 1 minute to lyse the red blood cells and resuspended in PBS. Single cells suspension was incubated with Fc Block TruStain FcX (Clone 93, Biolegend) in PBS with 5% normal rat serum and 5% normal mouse serum, and then stained with a cocktail Ab at predetermined concentration in staining buffer (PBS with 5% FBS and 25 mM HEPES). DAPI was used to exclude dead cells. The sorting was performed on BD Symphony S6 Cell Sorter.

### Single cell cDNA synthesis and sequencing library preparation

Tumors were collected 1 day after last dose of immunotherapy and total CD45 positive immune cells were sorted. Immediately after sorting, the sorted immune cells were fixed with Evercode Cell Fixation v2 (Parse Biosciences). the fixed cells were prepared using Evercode WT v2 kit (Parse Biosciences) according to manufacturer’s instructions. Input cell numbers and volumes were calculated using WT_100K_Sample_Loading_Table_V1.3.0 (Parse Biosciences). All the samples were processed together with Evercode WT v2 kit (Parse Biosciences) and 8 sub-libraries were generated. The quality of sub-libraries was checked by the Agilent Tape Station system (Agilent) and sequenced by Illumina NovaSeqX.

### Single-cell RNA-seq analysis

The raw FASTQ files were processed by spipe.v1.1.1(Parse Biosciences) using mm10 as a reference. Count matrices were imported to Seurat (v.4.3.0).(Hao et al., 2021) Cells with <50 detected genes or >10000 detected genes or > 7.5% mitochondrial genes were filtered-out from the dataset. We then used the standard Seurat pipeline by running NormalizeData(), FindVariableFeatures(selection.method =”vst”, nfeatures =4000), and ScaleData(). For the first round of clustering, Principal component analysis (PCA) was performed and the top 50 PCs with a resolution = 0.6 were applied. RunUMAP(dim=1:40, n.neighbours =30) was used to visualize the data. Cell types were annotated using SingleR (V1.7.1)(Aran et al., 2019) package with ImmGen data as reference. Neutrophils were subset, and standard pipeline was applied to neutrophil population by running FindVariableFeatures(selection.method =”vst”, nfeatures =4000), and ScaleData(). For clustering the neutrophils, Principal component analysis (PCA) was re-performed and the top 40 PCs with a resolution = 0.2 were applied. For visualization, RunUMAP(dim=1:50, n.neighbours =30) was used.

For RNA velocity analysis, spliced and unspliced count matrices were first obtained from spipe output and exported as h5ad files for subsequent input of scVelo. Velocity analysis was then performed with scVelo (0.2.5)(Bergen et al., 2020) using the default stochastic model and velocity vectors were projected into the UMAP generated from previous Seurat pipeline.

For pesudobulk analysis, cells were randomly grouped into 3 pseudo replicates. Gene counts were summarized and DESeq2 R package (v1.30.0)(Love et al., 2014) in R (v4.1.1) was used for differential gene analysis.

## Data and Software Availability

Raw and analyzed data is available at NCBI and the accession number for the single cell RNA sequencing reported in this paper are NCBI GEO: GSE278435,

## Acknowledgments

We thank C. Ciszewski at the Human Disease & Immune Discovery Core Facility at the UChicago for conducting FACS sorting; P. Faber (Genomics Core at the UChicago) for sequencing and raw data processing, Animal Resources Center at UChicago This study was supported by Y.M.’s Start-up fund and seed grants from The University of Chicago, grants to Y. M. from NIH (R00CA237859, R01CA285786 and R35GM150610), V Foundation, American Association for Cancer Research, and The Cancer Research Foundation.

## Author Contributions

Y.M., W.G. and D.L. conceptualized the study, designed the experiments, interpreted the data, and wrote the manuscript. W.G. and D.L. performed most experiments and analyzed data with the assistance from B.N. J.Q. provided LSL-Sox2 mice.

## Reference

Aran, D., A.P. Looney, L. Liu, E. Wu, V. Fong, A. Hsu, S. Chak, R.P. Naikawadi, P.J. Wolters, A.R. Abate, A.J. Butte, and M. Bhattacharya. 2019. Reference-based analysis of lung single-cell sequencing reveals a transitional profibrotic macrophage. Nat Immunol 20:163–172.

Bass, A.J., H. Watanabe, C.H. Mermel, S. Yu, S. Perner, R.G. Verhaak, S.Y. Kim, L. Wardwell, P. Tamayo, I. Gat-Viks, A.H. Ramos, M.S. Woo, B.A. Weir, G. Getz, R. Beroukhim, M. O’Kelly, A. Dutt, O. Rozenblatt-Rosen, P. Dziunycz, J. Komisarof, L.R. Chirieac, C.J. Lafargue, V. Scheble, T. Wilbertz, C. Ma, S. Rao, H. Nakagawa, D.B. Stairs, L. Lin, T.J. Giordano, P. Wagner, J.D. Minna, A.F. Gazdar, C.Q. Zhu, M.S. Brose, I. Cecconello, U. Ribeiro, Jr., S.K. Marie, O. Dahl, R.A. Shivdasani, M.S. Tsao, M.A. Rubin, K.K. Wong, A. Regev, W.C. Hahn, D.G. Beer, A.K. Rustgi, and M. Meyerson. 2009. SOX2 is an amplified lineage-survival oncogene in lung and esophageal squamous cell carcinomas. Nat Genet 41:1238–1242.

Benguigui, M., T.J. Cooper, P. Kalkar, S. Schif-Zuck, R. Halaban, A. Bacchiocchi, I. Kamer, A. Deo, B. Manobla, R. Menachem, J. Haj-Shomaly, A. Vorontsova, Z. Raviv, C. Buxbaum, P. Christopoulos, J. Bar, M. Lotem, M. Sznol, A. Ariel, S.S. Shen-Orr, and Y. Shaked. 2024. Interferon-stimulated neutrophils as a predictor of immunotherapy response. Cancer Cell 42:253–265 e212.

Bergen, V., M. Lange, S. Peidli, F.A. Wolf, and F.J. Theis. 2020. Generalizing RNA velocity to transient cell states through dynamical modeling. Nat Biotechnol 38:1408–1414.

Biswas, S.K., and A. Mantovani. 2010. Macrophage plasticity and interaction with lymphocyte subsets: cancer as a paradigm. Nat Immunol 11:889–896.

Boumahdi, S., G. Driessens, G. Lapouge, S. Rorive, D. Nassar, M. Le Mercier, B. Delatte, A. Caauwe, S. Lenglez, E. Nkusi, S. Brohee, I. Salmon, C. Dubois, V. del Marmol, F. Fuks, B. Beck, and C. Blanpain. 2014. SOX2 controls tumour initiation and cancer stem-cell functions in squamous-cell carcinoma. Nature 511:246–250.

Casanova-Acebes, M., E. Dalla, A.M. Leader, J. LeBerichel, J. Nikolic, B.M. Morales, M. Brown, C. Chang, L. Troncoso, S.T. Chen, A. Sastre-Perona, M.D. Park, A. Tabachnikova, M. Dhainaut, P. Hamon, B. Maier, C.M. Sawai, E. Agullo-Pascual, M. Schober, B.D. Brown, B. Reizis, T. Marron, E. Kenigsberg, C. Moussion, P. Benaroch, J.A. Aguirre-Ghiso, and M. Merad. 2021. Tissue-resident macrophages provide a pro-tumorigenic niche to early NSCLC cells. Nature 595:578–584.

Franklin, R.A., W. Liao, A. Sarkar, M.V. Kim, M.R. Bivona, K. Liu, E.G. Pamer, and M.O. Li. 2014. The cellular and molecular origin of tumor-associated macrophages. Science 344:921–925.

Ge, Y., and E. Fuchs. 2018. Stretching the limits: from homeostasis to stem cell plasticity in wound healing and cancer. Nat Rev Genet 19:311–325.

Ge, Y., N.C. Gomez, R.C. Adam, M. Nikolova, H. Yang, A. Verma, C.P. Lu, L. Polak, S. Yuan, O. Elemento, and E. Fuchs. 2017. Stem Cell Lineage Infidelity Drives Wound Repair and Cancer. Cell 169:636–650 e614.

Gonzales, K.A.U., L. Polak, I. Matos, M.T. Tierney, A. Gola, E. Wong, N.R. Infarinato, M. Nikolova, S. Luo, S. Liu, J.S.S. Novak, K. Lay, H.A. Pasolli, and E. Fuchs. 2021. Stem cells expand potency and alter tissue fitness by accumulating diverse epigenetic memories. Science 374:eabh2444.

Governa, V., K.G. de Oliveira, A. Bang-Rudenstam, S. Offer, M. Cerezo-Magana, J. Li, S. Beyer, M.C. Johansson, A.S. Mansson, C. Edvardsson, F. Durmo, E. Gustafsson, A. Boukredine, P. Jeannot, K. Schmidt, E. Gezelius, J.A. Menard, R. Garza, J. Jakobsson, T. de Neergaard, P.C. Sundgren, A.M. Tiihonen, H. Haapasalo, K.J. Rautajoki, P. Nordenfelt, A. Darabi, K. Forsberg-Nilsson, A. Pietras, H. Talbot, J. Bengzon, and M. Belting. 2024. Protumoral lipid droplet-loaded macrophages are enriched in human glioblastoma and can be therapeutically targeted. Sci Transl Med 16:eadk1168.

Gungabeesoon, J., N.A. Gort-Freitas, M. Kiss, E. Bolli, M. Messemaker, M. Siwicki, M. Hicham, R. Bill, P. Koch, C. Cianciaruso, F. Duval, C. Pfirschke, M. Mazzola, S. Peters, K. Homicsko, C. Garris, R. Weissleder, A.M. Klein, and M.J. Pittet. 2023. A neutrophil response linked to tumor control in immunotherapy. Cell 186:1448–1464 e1420.

Guo W, L.J., Huang X, Leon D, Good J, Nicholson B, Owusu-Ofori A, Izumchenko E, Rosenberg A, Agrawal N, Bertacchi B, Bolotin D, Gunzer M, Ballesteros I, Hidalgo A, Miao Y. . 2025. Tumor-Initiating Stem Cells Fine-tune the Plasticity of Neutrophils to Sculpt a Protective Niche. . bioRxiv 2025.04.04.647324

Guo, X., Y. Zhao, H. Yan, Y. Yang, S. Shen, X. Dai, X. Ji, F. Ji, X.G. Gong, L. Li, X. Bai, X.H. Feng, T. Liang, J. Ji, L. Chen, H. Wang, and B. Zhao. 2017. Single tumor-initiating cells evade immune clearance by recruiting type II macrophages. Genes Dev 31:247–259.

Han, C., V. Godfrey, Z. Liu, Y. Han, L. Liu, H. Peng, R.R. Weichselbaum, H. Zaki, and Y.X. Fu. 2021. The AIM2 and NLRP3 inflammasomes trigger IL-1-mediated antitumor effects during radiation. Sci Immunol 6:

Hao, Y., S. Hao, E. Andersen-Nissen, W.M. Mauck, 3rd, S. Zheng, A. Butler, M.J. Lee, A.J. Wilk, C. Darby, M. Zager, P. Hoffman, M. Stoeckius, E. Papalexi, E.P. Mimitou, J. Jain, A. Srivastava, T. Stuart, L.M. Fleming, B. Yeung, A.J. Rogers, J.M. McElrath, C.A. Blish, R. Gottardo, P. Smibert, and R. Satija. 2021. Integrated analysis of multimodal single-cell data. Cell 184:3573–3587 e3529.

Hegde, S., B. Giotti, B.Y. Soong, L. Halasz, J. Le Berichel, M.M. Schaefer, B. Kloeckner, R. Mattiuz, M.D. Park, A. Magen, A. Marks, M. Belabed, P. Hamon, T. Chin, L. Troncoso, J.J. Lee, K. Fan, D. Ahimovic, M.J. Bale, K. Nie, G. Chung, D. D’Souza, K. Angeliadis, S. Kim-Schulze, R.M. Flores, A.J. Kaufman, F. Ginhoux, J.D. Buenrostro, S.Z. Josefowicz, A.M. Tsankov, T.U. Marron, S. Ma, B.D. Brown, and M. Merad. 2025. Myeloid progenitor dysregulation fuels immunosuppressive macrophages in tumours. Nature

Hu, K.H., N.F. Kuhn, T. Courau, J. Tsui, B. Samad, P. Ha, J.R. Kratz, A.J. Combes, and M.F. Krummel. 2023. Transcriptional space-time mapping identifies concerted immune and stromal cell patterns and gene programs in wound healing and cancer. Cell Stem Cell 30:885–903 e810.

Huang, P.Y., E. Kandyba, A. Jabouille, J. Sjolund, A. Kumar, K. Halliwill, M. McCreery, R. DelRosario, H.C. Kang, C.E. Wong, J. Seibler, V. Beuger, M. Pellegrino, A. Sciambi, D.J. Eastburn, and A. Balmain. 2017. Lgr6 is a stem cell marker in mouse skin squamous cell carcinoma. Nat Genet 49:1624–1632.

Jing, L., Y. An, T. Cai, J. Xiang, B. Li, J. Guo, X. Ma, L. Wei, Y. Tian, X. Cheng, X. Chen, Z. Liu, J. Feng, F. Yang, X. Yan, and H. Duan. 2023. A subpopulation of CD146(+) macrophages enhances antitumor immunity by activating the NLRP3 inflammasome. Cell Mol Immunol 20:908–923.

Kartal, B., C.S. Garris, H.S. Kim, R.H. Kohler, J. Carrothers, E.A. Halabi, Y. Iwamoto, A.G. Goubet, Y. Xie, P. Wirapati, M.J. Pittet, and R. Weissleder. 2025. Targeted SPP1 Inhibition of Tumor-Associated Myeloid Cells Effectively Decreases Tumor Sizes. Adv Sci (Weinh) 12:e2410360.

Katzenelenbogen, Y., F. Sheban, A. Yalin, I. Yofe, D. Svetlichnyy, D.A. Jaitin, C. Bornstein, A. Moshe, H. Keren-Shaul, M. Cohen, S.Y. Wang, B. Li, E. David, T.M. Salame, A. Weiner, and I. Amit. 2020. Coupled scRNA-Seq and Intracellular Protein Activity Reveal an Immunosuppressive Role of TREM2 in Cancer. Cell 182:872–885 e819.

Kloosterman, D.J., J. Erbani, M. Boon, M. Farber, S.M. Handgraaf, M. Ando-Kuri, E. Sanchez-Lopez, B. Fontein, M. Mertz, M. Nieuwland, N.Q. Liu, G. Forn-Cuni, N.N. van der Wel, A.E. Grootemaat, L. Reinalda, S.I. van Kasteren, E. de Wit, B. Ruffell, E. Snaar-Jagalska, K. Petrecca, D. Brandsma, A. Kros, M. Giera, and L. Akkari. 2024. Macrophage-mediated myelin recycling fuels brain cancer malignancy. Cell 187:5336–5356 e5330.

Lapouge, G., K.K. Youssef, B. Vokaer, Y. Achouri, C. Michaux, P.A. Sotiropoulou, and C. Blanpain. 2011. Identifying the cellular origin of squamous skin tumors. Proc Natl Acad Sci U S A 108:7431–7436.

Liu, K., M. Jiang, Y. Lu, H. Chen, J. Sun, S. Wu, W.Y. Ku, H. Nakagawa, Y. Kita, S. Natsugoe, J.H. Peters, A. Rustgi, M.W. Onaitis, A. Kiernan, X. Chen, and J. Que. 2013. Sox2 cooperates with inflammation-mediated Stat3 activation in the malignant transformation of foregut basal progenitor cells. Cell Stem Cell 12:304–315.

Locati, M., G. Curtale, and A. Mantovani. 2020. Diversity, Mechanisms, and Significance of Macrophage Plasticity. Annu Rev Pathol 15:123–147.

Love, M.I., W. Huber, and S. Anders. 2014. Moderated estimation of fold change and dispersion for RNA-seq data with DESeq2. Genome Biol 15:550.

Lu, H., K.R. Clauser, W.L. Tam, J. Frose, X. Ye, E.N. Eaton, F. Reinhardt, V.S. Donnenberg, R. Bhargava, S.A. Carr, and R.A. Weinberg. 2014. A breast cancer stem cell niche supported by juxtacrine signalling from monocytes and macrophages. Nat Cell Biol 16:1105–1117.

Luan, J., C. Truong, A. Vuchkovska, W. Guo, J. Good, B. Liu, A. Gang, N. Infarinato, K. Stewart, L. Polak, H.A. Pasolli, E. Andretta, A.Y. Rudensky, E. Fuchs, and Y. Miao. 2024. CD80 on skin stem cells promotes local expansion of regulatory T cells upon injury to orchestrate repair within an inflammatory environment. Immunity

Marelli, G., N. Morina, F. Portale, M. Pandini, M. Iovino, G. Di Conza, P.C. Ho, and D. Di Mitri. 2022. Lipid-loaded macrophages as new therapeutic target in cancer. J Immunother Cancer 10:

Miao, Y., H. Yang, J. Levorse, S. Yuan, L. Polak, M. Sribour, B. Singh, M.D. Rosenblum, and E. Fuchs. 2019. Adaptive Immune Resistance Emerges from Tumor-Initiating Stem Cells. Cell 177:1172–1186 e1114.

Molgora, M., E. Esaulova, W. Vermi, J. Hou, Y. Chen, J. Luo, S. Brioschi, M. Bugatti, A.S. Omodei, B. Ricci, C. Fronick, S.K. Panda, Y. Takeuchi, M.M. Gubin, R. Faccio, M. Cella, S. Gilfillan, E.R. Unanue, M.N. Artyomov, R.D. Schreiber, and M. Colonna. 2020. TREM2 Modulation Remodels the Tumor Myeloid Landscape Enhancing Anti-PD-1 Immunotherapy. Cell 182:886–900 e817.

Mosser, D.M., and J.P. Edwards. 2008. Exploring the full spectrum of macrophage activation. Nat Rev Immunol 8:958–969.

Murray, P.J., J.E. Allen, S.K. Biswas, E.A. Fisher, D.W. Gilroy, S. Goerdt, S. Gordon, J.A. Hamilton, L.B. Ivashkiv, T. Lawrence, M. Locati, A. Mantovani, F.O. Martinez, J.L. Mege, D.M. Mosser, G. Natoli, J.P. Saeij, J.L. Schultze, K.A. Shirey, A. Sica, J. Suttles, I. Udalova, J.A. van Ginderachter, S.N. Vogel, and T.A. Wynn. 2014. Macrophage activation and polarization: nomenclature and experimental guidelines. Immunity 41:14–20.

Naik, S., and E. Fuchs. 2022. Inflammatory memory and tissue adaptation in sickness and in health. Nature 607:249–255.

Nassar, D., M. Latil, B. Boeckx, D. Lambrechts, and C. Blanpain. 2015. Genomic landscape of carcinogen-induced and genetically induced mouse skin squamous cell carcinoma. Nat Med 21:946–954.

Oshimori, N., D. Oristian, and E. Fuchs. 2015. TGF-beta promotes heterogeneity and drug resistance in squamous cell carcinoma. Cell 160:963–976.

Quintanilla, M., K. Brown, M. Ramsden, and A. Balmain. 1986. Carcinogen-specific mutation and amplification of Ha-ras during mouse skin carcinogenesis. Nature 322:78–80.

Raghavan, S., P. Mehta, Y. Xie, Y.L. Lei, and G. Mehta. 2019. Ovarian cancer stem cells and macrophages reciprocally interact through the WNT pathway to promote pro-tumoral and malignant phenotypes in 3D engineered microenvironments. J Immunother Cancer 7:190.

Ricci-Vitiani, L., D.G. Lombardi, E. Pilozzi, M. Biffoni, M. Todaro, C. Peschle, and R. De Maria. 2007. Identification and expansion of human colon-cancer-initiating cells. Nature 445:111–115.

Rosenberg, A.B., C.M. Roco, R.A. Muscat, A. Kuchina, P. Sample, Z. Yao, L.T. Graybuck, D.J. Peeler, S. Mukherjee, W. Chen, S.H. Pun, D.L. Sellers, B. Tasic, and G. Seelig. 2018. Single-cell profiling of the developing mouse brain and spinal cord with split-pool barcoding. Science 360:176–182.

Shen, C., J.H. Chen, H.R. Oh, and J.H. Park. 2021. Transcription factor SOX2 contributes to nonalcoholic fatty liver disease development by regulating the expression of the fatty acid transporter CD36. FEBS Lett 595:2493–2503.

Shen, C., H.R. Oh, Y.R. Park, S. Oh, and J.H. Park. 2025. Soluble DPP4 promotes hepatocyte lipid accumulation via SOX2-SCD1 signaling and counteracts DPP4 inhibition. Biochem Biophys Res Commun 756:151521.

Shook, B., E. Xiao, Y. Kumamoto, A. Iwasaki, and V. Horsley. 2016. CD301b+ Macrophages Are Essential for Effective Skin Wound Healing. J Invest Dermatol 136:1885–1891.

Shook, B.A., R.R. Wasko, G.C. Rivera-Gonzalez, E. Salazar-Gatzimas, F. Lopez-Giraldez, B.C. Dash, A.R. Munoz-Rojas, K.D. Aultman, R.K. Zwick, V. Lei, J.L. Arbiser, K. Miller-Jensen, D.A. Clark, H.C. Hsia, and V. Horsley. 2018. Myofibroblast proliferation and heterogeneity are supported by macrophages during skin repair. Science 362:

Singh, S.K., C. Hawkins, I.D. Clarke, J.A. Squire, J. Bayani, T. Hide, R.M. Henkelman, M.D. Cusimano, and P.B. Dirks. 2004. Identification of human brain tumour initiating cells. Nature 432:396–401.

Sun, L., T. Kees, A.S. Almeida, B. Liu, X.Y. He, D. Ng, X. Han, D.L. Spector, I.A. McNeish, P. Gimotty, S. Adams, and M. Egeblad. 2021. Activating a collaborative innate-adaptive immune response to control metastasis. Cancer Cell 39:1361–1374 e1369.

Sun, R., R. Han, C. McCornack, S. Khan, G.T. Tabor, Y. Chen, J. Hou, H. Jiang, K.M. Schoch, D.D. Mao, R. Cleary, A. Yang, Q. Liu, J. Luo, A. Petti, T.M. Miller, J.D. Ulrich, D.M. Holtzman, and A.H. Kim. 2023. TREM2 inhibition triggers antitumor cell activity of myeloid cells in glioblastoma. Sci Adv 9:eade3559.

Taniguchi, S., A. Elhance, A. Van Duzer, S. Kumar, J.J. Leitenberger, and N. Oshimori. 2020. Tumor-initiating cells establish an IL-33-TGF-beta niche signaling loop to promote cancer progression. Science 369:

Vannella, K.M., and T.A. Wynn. 2017. Mechanisms of Organ Injury and Repair by Macrophages. Annu Rev Physiol 79:593–617.

Vanner, R.J., M. Remke, M. Gallo, H.J. Selvadurai, F. Coutinho, L. Lee, M. Kushida, R. Head, S. Morrissy, X. Zhu, T. Aviv, V. Voisin, I.D. Clarke, Y. Li, A.J. Mungall, R.A. Moore, Y. Ma, S.J. Jones, M.A. Marra, D. Malkin, P.A. Northcott, M. Kool, S.M. Pfister, G. Bader, K. Hochedlinger, A. Korshunov, M.D. Taylor, and P.B. Dirks. 2014. Quiescent sox2(+) cells drive hierarchical growth and relapse in sonic hedgehog subgroup medulloblastoma. Cancer Cell 26:33–47.

Wang, Z., R. Dai, L. Kang, H. Yang, Z. Chen, J. He, L. Shu, Y. Zhong, Y. Zhang, Z. Hua, Y. Huang, Y. Jiang, J. Li, L. Xu, F. Lan, S.H. Lin, and J. Wong. 2025. SOX2 drives esophageal squamous carcinoma by reprogramming lipid metabolism and histone acetylation landscape. Nat Commun 16:8190.

Weiss, J.M., L.A. Ridnour, T. Back, S.P. Hussain, P. He, A.E. Maciag, L.K. Keefer, W.J. Murphy, C.C. Harris, D.A. Wink, and R.H. Wiltrout. 2010. Macrophage-dependent nitric oxide expression regulates tumor cell detachment and metastasis after IL-2/anti-CD40 immunotherapy. J Exp Med 207:2455–2467.

Wynn, T.A., A. Chawla, and J.W. Pollard. 2013. Macrophage biology in development, homeostasis and disease. Nature 496:445–455.

Wynn, T.A., and K.M. Vannella. 2016. Macrophages in Tissue Repair, Regeneration, and Fibrosis. Immunity 44:450–462.

Zhivaki, D., and J.C. Kagan. 2021. NLRP3 inflammasomes that induce antitumor immunity. Trends Immunol 42:575–589.

