## Supplemental Figure 1 for "Comparative single cell analysis of wound and cancer identifies the metabolic dialogues between tumor initiating stem cells and macrophages"

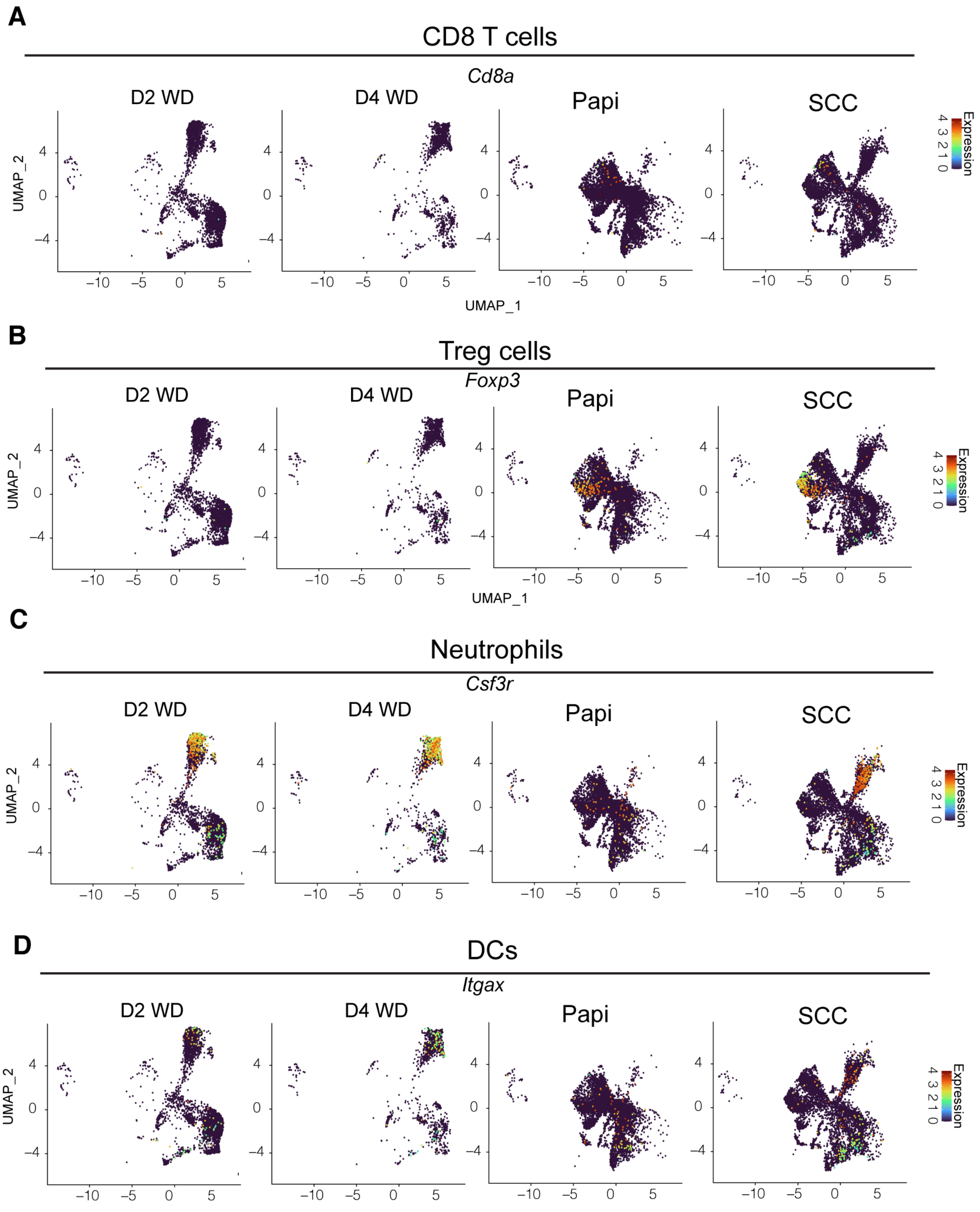
A
CD8 T cells
Cd8a
D2 WD
D4 WD
Papi
SCC
UMAP_2
UMAP_1
Expression
B
Treg cells
Foxp3
D2 WD
D4 WD
Papi
SCC
UMAP_2
UMAP_1
Expression
C
Neutrophils
Csf3r
D2 WD
D4 WD
Papi
SCC
UMAP_2
Expression
D
DCs
Itgax
D2 WD
D4 WD
Papi
SCC
UMAP_2
Expression

**Figure S1. Global immune landscape in wound and cancer. Related to Figure 1.**
**a** to **d**. UMAP showing the expression of signature genes used for identifying (**a**) CD8 T cells, (**b**) Treg cells, (**c**) Neutrophils, and (**d**) DCs.
